# Modulation of brain activity by psycholinguistic information during naturalistic speech comprehension and production

**DOI:** 10.1101/2022.03.07.483336

**Authors:** Wei Wu, Matías Morales, Tanvi Patel, Martin J. Pickering, Paul Hoffman

## Abstract

Language processing requires the integration of diverse sources of information across multiple levels of processing. A range of psycholinguistic properties have been documented in previous studies as having influence on brain activation during language processing. However, most of those studies have used factorial designs to probe the effect of one or two individual properties using highly controlled stimuli and experimental paradigms. Little is known about the neural correlates of psycholinguistic properties in more naturalistic discourse, especially during language production. The aim of our study is to explore the above issues in a rich fMRI dataset in which participants both listened to recorded passages of discourse and produced their own narrative discourse in response to prompts. Specifically, we measured 13 psycholinguistic properties of the discourse comprehended or produced by the participants, and we used principal components analysis (PCA) to address covariation in these properties and extract a smaller set of latent language characteristics. These latent components indexed vocabulary complexity, sensory-motor and emotional language content, discourse coherence and speech quantity. A parametric approach was adopted to study the effects of these psycholinguistic variables on brain activation during comprehension and production. We found that the pattern of effects across the cortex was somewhat convergent across comprehension and production. However, the degree of convergence varied across language properties, being strongest for the component indexing sensory-motor language content. We report the full, unthresholded effect maps for each psycholinguistic variable, as well as mapping how these effects change along a large-scale cortical gradient of brain function. We believe that our findings provide a valuable starting point for future, confirmatory studies of discourse processing.

## 1. Introduction

Human brains possess the ability to understand and produce language with apparent ease, which provides the foundation for social interaction and communication in our everyday lives. Although language processing has always been a popular topic of study in cognitive science, linguistics and other fields, the neural substrate supporting language processing is still a puzzle, particularly at the level of complex discourse. Processing of language is a multifaceted process that requires the integration of diverse sources of information across multiple levels of processing. Each of these levels engages distinct brain networks, which are in turn responsive to different properties of speech. At the level of auditory-phonological perception, for example, classic studies have demonstrated linear response increases in the auditory system in relation to the rate of speech input (Dhankhar et al., 1997; Price et al., 1992). At the semantic level, embodiment theories predict the engagement of specific sensory-motor association cortices when people process language that relates to different types of sensory-motor experience (Barsalou, 2008; Glenberg & Gallese, 2012). And at the discourse level, executive control regions are thought to play an important role in regulating the topic of speech and have been implicated in the maintenance of coherence in narrative production tasks (Ash et al., 2013; Hoffman, 2019; Marini & Andreetta, 2016). Consequentially, in order to build models of the neural basis of language, it is critical to understand how activation across the brain varies as a function of different properties of language across processing levels.

fMRI investigations of the effect of psycholinguistic properties on activation commonly use factorial manipulations of one or two individual properties in highly controlled experiments (e.g., Binder, Westbury, McKiernan, Possing, & Medler, 2005; Hoffman, Binney, & Lambon Ralph, 2015; Skipper & Olson, 2014; Xiaosha Wang, Wang, & Bi, 2019). For example, to investigate which regions are sensitive to the concreteness of words, researchers will typically employ two sets of stimuli which differ in concreteness but are matched for other properties that the researchers deem important (for meta-analyses of many such studies, see Bucur & Papagno, 2021; J. Wang, Conder, Blitzer, & Shinkareva, 2010). Factorial designs such as these are ideal for ensuring stimulus control and have led to many important discoveries in the neuroscience of language. They are, however, subject to some limitations. By adopting a binary assignment to two conditions, factorial analyses are insensitive to continuous variation in the property of interest within each condition. In addition, factorial designs frequently do not exploit the full range of values available for continuous variables. To combat this, an alternative approach is to investigate the neural response to the property of interest in a parametric fashion and some studies have done this with great success (Graves, Desai, Humphries, Seidenberg, & Binder, 2010; Hauk, Davis, & Pulvermuller, 2008; Mummery, Ashburner, Scott, & Wise, 1999; Price et al., 1992; Wise et al., 2000).

In recent years, researchers have often moved beyond univariate analysis of single-voxel responses, in favour of sophisticated multivariate pattern analyses (MVPA) (Kriegeskorte & Kievit, 2013; Kriegeskorte, Mur, & Bandettini, 2008; Norman, Polyn, Detre, & Haxby, 2006). These techniques have been extremely valuable in identifying the organisational principles that support coding of different semantic and linguistic categories in the brain (Bruffaerts et al., 2019; Caucheteux & King, 2022; Frankland & Greene, 2015; Gao et al., 2021; Meersmans et al., 2021; Xu et al., 2018). However, they are less suited to identifying how stimulus properties influence the *degree* to which a brain region is activated. Classifiers trained on multivoxel data from a region can successfully discriminate between two experimental conditions, even when there is no mean activation difference between the conditions at the group level (for discussion, see Coutanche, 2013; Jimura & Poldrack, 2012; Mur, Bandettini, & Kriegeskorte, 2009). Furthermore, MVPA statistics are typically insensitive to the direction of an effect, meaning that significant multivariate effects can be found when the direction of the mean activation difference varies substantially between participants (Todd, Nystrom, & Cohen, 2013). For example, a classifier could show above-chance decoding for concrete vs. abstract words in a region, even if half of the participants showed greater mean activation for concrete words and the remainder showed the opposite effect. For these reasons, univariate analyses remain valuable when investigating how specific stimulus properties affect neural engagement.

Irrespective of the analysis approach, neurolinguistic studies have traditionally used single words or highly controlled simple sentences to probe language functions. A number of researchers have argued that these stimuli are not representative of language usage in everyday life and that a more naturalistic approach is essential to understand the neural correlates of language processing “in the wild” (Hamilton & Huth, 2020; Hasson & Honey, 2012; Nastase, Goldstein, & Hasson, 2020; Willems, Nastase, & Milivojevic, 2020). Although the aim of all studies is to generate conclusions that can be generalised to how language is used in real life, some effects observed in artificial experimental settings might not carry over to more naturalistic speech. At present, it is often unclear which psycholinguistic property effects observed at the single-word or sentence level are present when people process more natural language passages. For instance, theories of concreteness effects state that abstract words are harder to comprehend than concrete words because they are associated with more possible contexts (Schwanenflugel & Shoben, 1983). The additional processing demands associated with selecting an appropriate contextual interpretation has been proposed as an explanation for higher activation in prefrontal cortex for more abstract words (Bucur & Papagno, 2021; Hoffman, Binney, et al., 2015; J. Wang et al., 2010). Evidence for these effects comes primarily from studies of single words or sentences presented out of context. Such effects might not occur in more natural discourse, where a rich prior context is available to constrain semantic processing.

In attempts to increase ecological validity, an increasing number of researchers have used naturalistic language stimuli, such as stories, to investigate the neurobiology of language (e.g., Gwilliams, King, Marantz, & Poeppel, 2020; Huth, de Heer, Griffiths, Theunissen, & Gallant, 2016; Wehbe et al., 2021; Zhang, Han, Worth, & Liu, 2020). The predictors of neural activity in such studies vary. Some studies have used the response time course in one subjects brain to predict that in other individuals listening to the same story (i.e., inter-subject correlation analysis, Nastase, Gazzola, Hasson, & Keysers, 2019). These studies indicate that activity in a wide set of regions is synchronised across listeners during comprehension, and that this synchronisation is disrupted when people understand narratives in different ways (Nguyen, Vanderwal, & Hasson, 2019; Yeshurun et al., 2017). Others have used computational linguistic models of lexical-semantic content as predictors of neural activity, revealing the brain regions that encode different categories of semantic knowledge (Dehghani et al., 2017; Deniz, Nunez-Elizalde, Huth, & Gallant, 2019; Huth et al., 2016; Wehbe et al., 2014; Zhang et al., 2020). The use of traditional psycholinguistic variables, such as word frequency and concreteness, to predict activation is less common. Some studies have investigated this with story reading paradigms. For example, Desai and colleagues have combined fMRI and eye tracking to examine the effects of noun manipulability and word frequency in the brain during the reading of whole text paragraphs (Desai, Choi, & Henderson, 2020; Desai, Choi, Lai, & Henderson, 2016). Another group of researchers have investigated the neural correlates of various psycholinguistic properties of single words during the reading of rapidly presented narrative text (200–300ms /word) (Yarkoni, Speer, Balota, McAvoy, & Zacks, 2008). However, investigations in spoken comprehension are rare and limited to investigations of the effects of lexical surprisal on activation (Russo et al., 2020; Willems, Frank, Nijhof, Hagoort, & van den Bosch, 2016).

Although there has been a recent shift to investigating *comprehension* with more natural stimuli, far fewer studies have explored the neural correlates of language properties during discourse *production.* Almost all neuroimaging studies that have probed psycholinguistic property effects have done so in receptive (comprehension) tasks and little is known about whether similar effects occur during language production. Many theories of language processing posit shared processes and representations for comprehension and production, at the semantic and conceptual levels (Dell & Chang, 2014; Garrod & Pickering, 2004; Hagoort, 2013; Hickok & Poeppel, 2007; Kintsch & Van Dijk, 1978; Levelt, Roelofs, & Meyer, 1999; Pickering & Garrod, 2021). These theories predict that neural correlations with lexical-semantic properties should be similar irrespective of whether participants are listening to speech or producing their own utterances. However, the degree to which speech properties have similar influences on speaking and listening is largely unknown, because few studies have systematically investigated production, and fewer still have directly compared comprehension and production in the same individuals. Although some previous studies have examined the neural activity coupling across the speaker’s and listener’s brains during production and comprehension (Heidlmayr, Weber, Takashima, & Hagoort, 2020; Jiang et al., 2012; Liu et al., 2019; Nguyen et al., 2022; Silbert, Honey, Simony, Poeppel, & Hasson, 2014; Stephens, Silbert, & Hasson, 2010), those studies do not compare psycholinguistic property effects during speaking and listening in the same individuals.

We have recently acquired an fMRI dataset of naturalistic discourse comprehension and production that allows the above issues to be explored. In our study, during scanning, the same group of participants listened to excerpts of naturalistic speech on everyday topics as well as producing their own discourse in response to topic prompts (e.g., Describe how you would make a cup of tea or coffee). We originally used these data to investigate the correlates of discourse coherence on activation during comprehension and production (Morales, Patel, Tamm, Pickering, & Hoffman, in press). In the present exploratory study, we investigated the neural correlates of a wider range of psycholinguistic properties, comparing their effects during comprehension and production. Specifically, *we* used a parametric approach to investigate how neural activation is affected by fluctuations in the psycholinguistic properties of natural speech, characterising our results at different neural levels (i.e., whole-brain and network levels).

We used principal components analysis (PCA) to address covariation in language properties and to generate a small number of latent underlying psycholinguistic factors. All studies in the field of language processing have to deal with the fact that psycholinguistic properties are inter-correlated, such that the effects of one variable can be confounded by others. Later acquired words, for example, tend to be longer, lower in frequency and more abstract than those acquired early in development. In experimental paradigms, this issue can be addressed by varying one property while controlling for other correlated properties (e.g., Price et al., 1992; Xiaosha Wang et al., 2019; Wise et al., 2000) or by selecting stimuli such that the properties under investigation are uncorrelated (Graves et al., 2010). However, these methods cannot be adopted with naturalistic data, where researchers are at the mercy of statistical regularities present in the language as a whole. More importantly, the correlations between individual psycholinguistic properties may be a consequence of more primitive or basic qualities that underpin the structure of language. For example, the degree to which words evoke visual and haptic experiences are positively correlated with each other and with concreteness ratings (Connell & Lynott, 2012), presumably because all of these measures are influenced by the degree to which words refer to tangible objects.

The most commonly used and well-understood strategy to solve the above problems is to use PCA to extract a smaller set of latent variables that capture the underlying variance among a set of measures (for examples of the use of this approach in psycholinguistics studies, see Baayen, 2010; Baayen, Feldman, & Schreuder, 2006; Davies, Barbon, & Cuetos, 2013; Troche, Crutch, & Reilly, 2017). PCA applied to speech properties has been used successfully in previous studies of discourse and aphasia (e.g., Alyahya, Halai, Conroy, & Lambon Ralph, 2020a; Binder et al., 2016; Hoffman, Loginova, & Russell, 2018; Sajjadi, Patterson, Arnold, Watson, & Nestor, 2012). In particular, this strategy has been found to be useful in structural MRI studies to identify the neural correlates of different aspects of speech production in aphasic patients (Alyahya, Halai, Conroy, & Lambon Ralph, 2020b; Mirman et al., 2015). The use of PCA or other related data reduction methods is also implicit in fMRI studies which use computational models of natural language processing (NLP) to predict brain activation (Dehghani et al., 2017; Deniz et al., 2019; Huth et al., 2016; Pereira et al., 2018; Wehbe et al., 2014; Zhang et al., 2020). This is because such models use data reduction methods (sometimes through the use of deep neural networks) to extract latent components of word meaning from statistical patterns of word cooccurrences in large language corpora (Landauer & Dumais, 1997; Mikolov, Chen, Corrado, & Dean, 2013; Pennington, Socher, & Manning, 2014).

Although the general benefits of PCA are well-established in language research, the technique has rarely been applied to traditional psycholinguistic measures (as opposed to NLP language models) to generate predictors for fMRI analysis. We are aware of two previous studies. Hauk et al. (2008) applied PCA to a set of 21 word-level properties and used the resulting components to predict activation in a single-word reading task. More recently, Fernandino et al. (2016) used PCA on a set of 5 variables indexing sensory-motor experiences associated with words, in an fMRI study based on single-word semantic judgements. However, they did not investigate the neural correlates of the individual PCA components. In the current study, we examined whether PCA could be used to explore the core structure that underlies psycholinguistic properties in more naturalistic discourse comprehension and production tasks. We then investigated how neural activation correlated with variations in the latent psycholinguistic components we obtained.

In summary, in the present study, we aimed to directly compare the neural correlates of psycholinguistic effects in discourse during comprehension and production in a naturalistic task. Our approach was (1) to compute a range of language statistics from the discourse samples which participants listened to or produced in our study; (2) to perform PCA to investigate the underlying factors that emerged from the set of measures; (3) to explore how these underlying factors were correlated with neural activation at different levels. Rather than test specific hypotheses about which brain regions correlate with which aspects of language, our intention in this exploratory report is to present the full, unthresholded effect maps for each component of language and to map how these patterns change along a large-scale cortical gradient of brain function (Margulies et al., 2016). We hope that this exploratory investigation of naturalistic discourse processing will provide a useful starting point for future, targeted investigations of how specific regions contribute to discourse.

## 2. Materials and Methods

### 2.1. Participants

Twenty-five adult participants (21 females, mean age = 24 years, SD = 4.4 years, range = 18-35 years) were recruited from the University of Edinburgh and participated in the study in exchange for payment. Sample size was determined by the resources available to complete the study. No participants or runs were excluded from analysis. All participants were native speakers of English, righthanded based on the Edinburgh Handedness Inventory (Oldfield, 1971) and reported to be in good health with no history of neurological or psychiatric illness. The study was approved by the Psychology Research Ethics Committee of the University of Edinburgh and all participants gave informed consent.

### 2.2. Materials

In the comprehension and production tasks, discourse was related to 12 prompts that asked about common semantic knowledge on particular topics (e.g., How would you prepare to go on holiday? see Supplementary Table S1 for a complete list of prompts). Speech comprehension topics were different from those used in the production task, to avoid priming participants’ production responses with information presented in the comprehension trials. For the comprehension task, we selected 24 samples of speech (half of them were highly coherent and half of them were less coherent passages) discussing the 12 different topics from a corpus of responses provided by participants in a previous behavioural study, which were further split into two sets of different stimuli and each presented to half of the participants (Hoffman et al., 2018). All comprehension speech passages were recorded by the same male native English speaker and their duration was 50 s each. Baseline conditions involved either listening to (comprehension condition) or reciting (production condition) of the English nursery rhyme, Humpty Dumpty. Thus the baseline condition involved grammatically well-formed continuous speech, but without the requirement to understand or generate novel utterances. The baseline conditions were part of the dataset but not used in analyses in the current study.

### 2.3. Procedure

There were two production and two comprehension runs for each participant. Half of the participants began the experiment with a production run and the other half began with a comprehension run. Each run consisted of six discourse trials and five baseline trials, the order of which was fully randomised. The structure of a single trial is shown in Figure 1A. Specifically, for the discourse conditions, trials started with the presentation of a written prompt for 6 s. Participants were asked to prepare to listen/speak during this period and to start listening/speaking when a green circle replaced the prompt in the centre of screen. Participants were instructed to listen/speak for 50 s, after which a red cross would replace the green circle. At this point participants were instructed to wait for the next prompt to appear on screen. The procedure for the baseline conditions was the same as the discourse conditions, except that the participants were asked to listen (comprehension task) to or to recite (production task) the Humpty Dumpty rhyme for 10 s. All trials were presented with an interstimulus interval jittered between 3 s and 7 s (5 s on average) and each scanning run was approximately 8 min in total.

**Figure 1.**
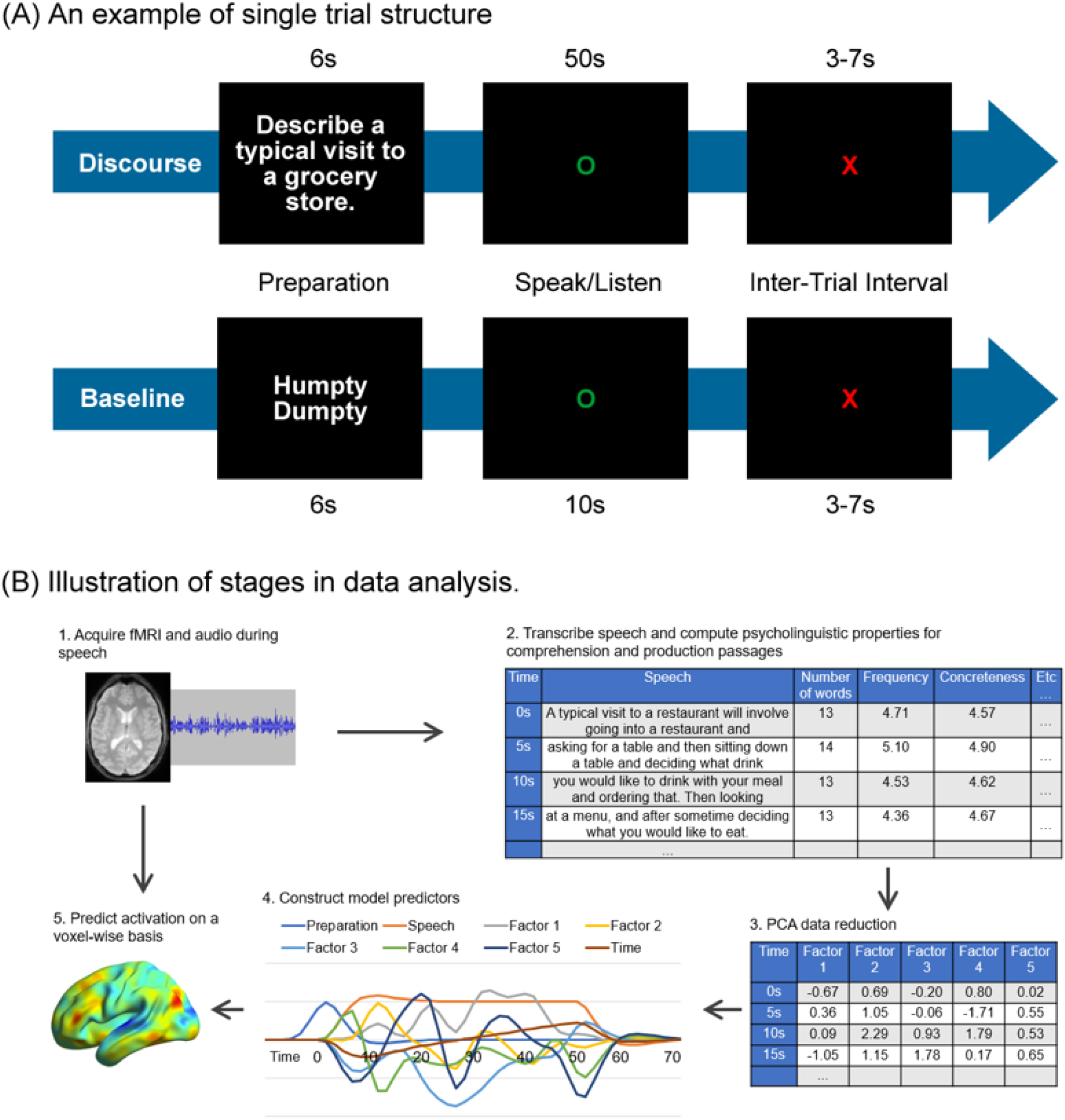
(A) Illustration of a single trial of the discourse and baseline tasks. (B) Stages in data analysis.

**Figure 2.**
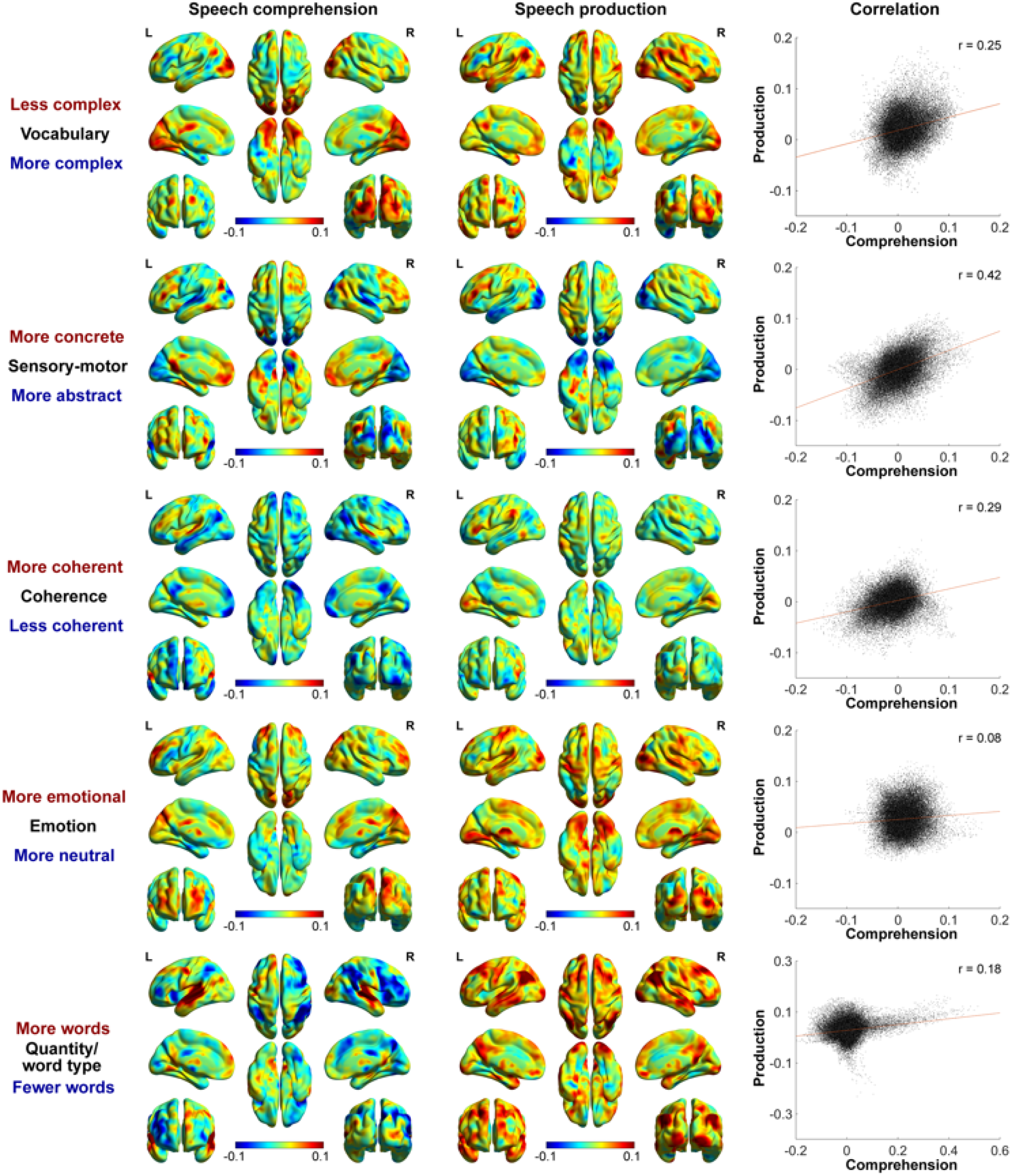
Unthresholded maps of psycholinguistic PC effects (beta values) on speech comprehension and speech production, and voxelwise correlations between tasks.

Before scanning, participants were presented with two training trials to familiarise them with the two tasks. They were also informed that they would receive a memory test regarding the comprehension runs after scanning to increase their motivation and attention. In this memory test, for each topic presented during speech comprehension runs, three statements were presented and one of them was taken from the actual comprehension passage. Participants were asked to choose the statement that they remembered hearing. Additionally, an audibility scale of 1 (inaudible) - 7 (perfectly audible) was rated to ensure participants could hear the audio. The results showed that participants correctly responded to most of the questions in the memory test (10 out of 12 on average, SD = 1.6; one-sample t-test comparing with chance performance: t = 31.21, *P* < 0.001) and rated the recordings in the comprehension task as highly audible (mean = 5.5, SD = 1.0).

### 2.4. Processing of speech samples

The overall analysis strategy is shown in Figure 1B. For the speech production runs, responses to each prompt were digitally recorded with an MRI-compatible microphone and processed with noise cancellation software (Cusack, Cumming, Bor, Norris, & Lyzenga, 2005) to reduce noise from the scanner. They were then transcribed and split into 5-second blocks. A block length of 5 s was chosen for two reasons. First, changes in some of our psycholinguistic properties occur only over relatively long timescales of discourse. For example, a shift in discourse away from its original topic (i.e., a change in global coherence) typically occurs over the course of at least one or two sentences. Second, a block length of 5 s is well matched to the temporal resolution of the BOLD signal. However, we conducted a control analysis where PCA was performed on data using windows of 10-second duration and this provided very similar results to those reported in the main analysis (see Supplementary Table S2).

We then calculated a number of psycholinguistic properties for each block of speech. We chose 13 variables that together index key aspects of lexical quality, semantic content, discourse structure and syntactic structure. While there are many other variables that we could potentially have included, we aimed to use the variables most commonly investigated in previous fMRI studies, such as word length, frequency and concreteness. We did not use vector-based semantic representations derived from computational models of language (Landauer & Dumais, 1997; Mikolov et al., 2013; Pennington et al., 2014), as their ability to predict activation has been investigated extensively in other work (Dehghani et al., 2017; Deniz et al., 2019; Huth et al., 2016; Pereira et al., 2018; Wehbe et al., 2014; Zhang et al., 2020).

Most of the variables were calculated at the lexical level by averaging the relevant measures over the nouns produced in the block (where words straddled a block boundary, their properties were counted in the block that contained the larger portion of the word). We restricted analysis of lexical properties to nouns to ensure that measures were comparable across blocks that contained different classes of word. Since different parts of speech vary systematically in their properties (e.g., verbs and adjectives tend to be less concrete than nouns; Bird, Franklin, & Howard, 2001), including all parts of speech could potentially confound lexical-semantic characteristics with syntactic structure and word class. However, we conducted a control analysis including all open-class words which provided very similar results (see Supplementary Table S3). We submitted transcripts to the Stanford Log-linear Part-of-Speech Tagger v3.8 for automated part-of-speech tagging (Toutanova, Klein, Manning, & Singer, 2003) in order to identify nouns. At the lexical level, the following psycholinguistic properties were then measured and averaged over words to give values for each block of speech:

#### Mean noun frequency

Log-transformed frequencies (Zipf) in the SUBTLEX-UK database (van Heuven, Mandera, Keuleers, & Brysbaert, 2014) were obtained for all words tagged as nouns.

#### Mean noun concreteness

Concreteness ratings of nouns were obtained from Brysbaert, Warriner, and Kuperman (2014).

#### Mean noun age of acquisition (AoA)

Estimates of AoA were obtained from the norms of Kuperman, Stadthagen-Gonzalez, and Brysbaert (2012).

#### Mean noun number of phonemes

The length of all nouns (in phonemes) was also calculated with the norms of Kuperman et al. (2012).

#### Mean noun semantic diversity (SemD)

SemD values for nouns were obtained from Hoffman, Lambon Ralph, and Rogers (2013). SemD is a measure of variability in the contextual usage of words. Words with high SemD values are used in a wide variety of contexts and thus have more variable and less well-specified meanings.

#### Mean noun perception strength

The perception strength (Minkowski 3 measure) of all nouns was obtained from the norms of Lynott, Connell, Brysbaert, Brand, and Carney (2020). This property was measured with behavioural ratings in which participants were asked to rate to what extent they experienced the concepts by six perceptual modalities (touch, hearing, smell, taste, vision, and interoception), and these single modality ratings were then combined to compute the perception strength.

#### Mean noun action strength

The action strength (Minkowski 3 measure) was obtained from the norms of Lynott et al. (2020). This property was measured with behavioural ratings in which participants were asked to rate to what extent they experienced the concepts by performing an action with five body effectors (mouth/throat, hand/arm, foot/leg, head excluding mouth/throat, and torso), and the ratings were combined to compute the action strength.

#### Mean noun valence

Valence ratings for nouns were obtained from Warriner, Kuperman, and Brysbaert (2013). As we were interested in the comparison of neutral versus highly valenced words (but not positive versus negative valence), we subtracted median value of the scale from the averaged valence ratings and transformed the results into absolute values.

#### Mean noun arousal

Arousal ratings in the norms of Warriner et al. (2013) were obtained for nouns in each block.

In addition, we measured the following properties at the level of the discourse:

#### Number of words

the total number of words produced in the 5-second speech block.

#### Proportion closed-class words

Closed-class words are words that play primarily a functional or syntactic role in language, as opposed to open-class words which carry semantic meaning. We classified nouns, verbs and adjectives and some adverbs (classified by the Stanford Part-of-Speech Tagger) as open-class and all other words (including pronouns, numbers, prepositions, conjunctions, determiners, auxiliaries and some adverbs) as closed-class. We then calculated the proportion of words in each block that were closed-class.

#### Local coherence

Local coherence refers to the degree to which adjoining utterances in speech are meaningfully related to one another. A measure of local coherence was computed using the same computational methods as Hoffman et al. (2018). Latent semantic analysis (LSA) was used to compare the semantic content of each 20-word passage of speech with the speech from the previous 20 words. High scores indicate a strong semantic relationship between adjoining passages of speech, whereas low scores indicate a shift in topic. A moving window approach was used to generate a value for the local coherence at each point in the discourse and the values for the words in each block were averaged.

#### Global coherence

Global coherence refers to the degree to which utterances were meaningfully related to the topic being probed. This property was measured using the method first described in Hoffman et al. (2018), by comparing the LSA semantic representation of each participants response to each prompt with a group-average prototype LSA vector that represented typical discourse on the topic. Higher values indicate utterances that are more closely related to typical discourse on the topic. This measure was again calculated over passages of 20 words, using a moving-window approach, with the value from each window was assigned to the final word in the window. The global coherence was included in the current study because of its close relationship with semantic control and executive function during speech processing (Hoffman, 2019; Hoffman et al., 2018; Morales et al., in press). This measure is the same as that used in our previous study (Morales et al., in press).

The above psycholinguistic properties were also quantified for the speech samples presented in the comprehension task. Thus, we obtained a comprehensive set of speech properties for each of the 3000 5-s blocks of speech recorded in the production task (10 blocks x 12 topics x 25 participants) and for each of the 240 blocks of speech presented to participants in the comprehension task (10 blocks x 12 topics x 2 versions of each topic).

### 2.5. Processing of psycholinguistic characteristics of speech

Considering that some of the psycholinguistic properties of interest may be inter-correlated with each other, a series of analyses were conducted to reveal the relationships among different properties and combine these properties into a comprehensive measure. The data for these analyses comprised the speech measures for the 3240 blocks of speech in the study (combining comprehension and production). Specifically, we first computed the Pearson correlation between each pair of psycholinguistic properties to identify covariations between different psycholinguistic properties. To further explore the structure among speech characteristics and to generate comprehensive measures for all psycholinguistic properties, a PCA was performed, which resulted in the extraction of five latent factors (which were the only factors with eigenvalues greater than one and together explained 68% of the variance within the set of properties). The factors were promax rotated to aid interpretation and to reduce correlations between the latent variables. Each block’s scores on these psycholinguistic principal components (PCs) were later used as predictors of neural activity in our main analyses (see Supplementary Figure S1 for the distribution of PC scores). There are a range of other data reduction approaches that could be applied to these data. We opted to use PCA because it is widely used in psycholinguistic research (see Introduction) and because of specific drawbacks associated with other approaches. For example, non-negative matrix factorisation imposes non-negativity constraints and generates factors that are not necessarily orthogonal to one another, which make this method tend to group only positively correlated variables together and ignore negative relationship that likely reflects a single underlying latent factor.

### 2.6. Image acquisition and processing

Images were acquired on a 3T Siemens Prisma scanner with a 32-channel head coil. For the functional images, the multi-echo EPI sequence included 46 slices covering the whole brain with echo time (TE) at 13 ms, 31 ms and 48 ms, repetition time (TR) = 1.7 s, flip angle = 73°, 80 × 80 matrix, reconstructed in-plane resolution = 3 mm × 3 mm, slice thickness = 3.0 mm (no slice gap) and multiband factor = 2. In total, four runs of 281 volumes (477.7 s) were acquired. A high-resolution T1-weighted structural image was also acquired for each participant using an MP-RAGE sequence with 1 mm isotropic voxels, TR = 2.5 s, TE = 4.6 ms. To minimize the impact of speech-related head movements and signal drop out in the ventral temporal regions (Kundu et al., 2017), the study employed a whole-brain multi-echo acquisition protocol, in which data were simultaneously acquired at 3 TEs. Data from the three echo series were weighted and combined, and the resulting time-series were denoised using independent components analysis (ICA).

Images were pre-processed and analysed using SPM12 and the TE-Dependent Analysis Toolbox 0.0.7 (Tedana) (Kundu et al., 2013; Kundu, Inati, Evans, Luh, & Bandettini, 2012). Estimates of head motion were obtained using the first BOLD echo series. Slice-timing correction was carried out and images were then realigned using the previously obtained motion estimates. Tedana was used to combine the three echo series into a single-time series and to divide the data into components classified as either BOLD-signal or noise-related based on their patterns of signal decay over increasing TEs (Kundu et al., 2017). Components classified as noise were discarded. After that, images were unwarped with a B0 fieldmap to correct for irregularities in the scanner’s magnetic field. Finally, functional images were spatially normalised to MNI space using SPM’s DARTEL tool (Ashburner, 2007) and were smoothed with a kernel of 8mm FWHM.

Data in our study were treated with a high-pass filter with a cut-off of 128 s and the four experimental runs (two comprehension and two production runs) were analysed using a single general linear model. Four speech periods were modelled as different event types: discourse comprehension, baseline comprehension, discourse production, and baseline production. Discourse periods were modelled as a series of concatenated 5-s blocks. This allowed us to include parametric modulators that coded the psycholinguistic PCs/properties of speech in each 5-s block. In our main analyses, the PC scores of the 5 psycholinguistic PCs, calculated as described earlier, were included in the model as modulators for each run. We also included time within each discourse period as another modulator to exclude the potential influences of time on effects of psycholinguistic information (e.g., later stages of a speech tend to possess lower coherence, see Hoffman, 2019). Modulators were mean-centred for each run. Additional regressors modelled the preparation periods for discourse and baseline in each task. Covariates consisted of six motion parameters and their first-order derivatives. In addition to the PC-based model, we also conducted parametric modulation analysis for each individual psycholinguistic property separately by including one property at a time in the GLM. This provided data on the raw property effects without considering relationships among different psycholinguistic properties. We include these results as supplementary analyses.

### 2.7. Analyses

After estimation of the first-level models, we submitted the individual-level beta maps of each modulator to second-level group analyses. In keeping with the exploratory nature of the investigation, our analyses focus on characterising the effect sizes of each PC in different parts of the brain, rather than testing specific hypotheses.

#### Whole-brain level

In this section of analyses, we investigated the effects of psycholinguistic PCs/properties in areas across the whole brain. Individual-level beta maps of each psycholinguistic modulator were averaged across all participants. In line with the guidelines for *Cortex’s* Exploratory Reports format, we present the unthresholded group-level beta maps, which provide estimates of the effect sizes of different psycholinguistic PCs/properties on activation throughout the brain. The brain activation maps were visualized with the BrainNet Viewer (http://www.nitrc.org/projects/bnv/) (Xia, Wang, & He, 2013). We did not threshold results based on tests of statistical significance (again in line with guidelines for Exploratory Reports) as our aim was to provide visualisations of the overall pattern of effects across the whole brain. To investigate the similarity of the beta maps in comprehension and production for each psycholinguistic PC, we extracted the beta values for comprehension and production in each voxel and computed voxelwise correlations between the two tasks.

#### Network level

We also investigated the effect of psycholinguistic information on activation at network level. Cortical networks are often identified in a discrete fashion, e.g., by using connectivity patterns in resting-state fMRI data to segregate the cortex into a set of distinct networks (e.g., Yeo et al., 2011). Such approaches assume hard boundaries between networks. Here, however, we used a different approach based on the assumption that function varies in a graded fashion as one moves across the cortical surface. We used the principal connectivity gradient described in Margulies et al. (2016). Margulies et al. mapped the organisation of the cortex along a single continuous gradient, such that regions of the brain that shared similar patterns of functional connectivity were located at similar points on the gradient (shown in Figure 3A). At one end of this spectrum lie the sensorimotor cortices, which show strong functional connectivity with one another. At the other end lie regions associated with the default-mode network (DMN), whose activity is correlated with one another but is anticorrelated with sensorimotor systems. Regions situated between the two extreme ends of the principal gradient include the inferior frontal sulcus, the intraparietal sulcus, and the inferior temporal sulcus, constituting heteromodal integration and higher-order cognitive regions (e.g., attention and executive areas). It has been proposed that this spectrum represents a functional hierarchy in the cortex, ranging from regions implicated in external, stimulus-driven processing to those engaged by internally-generated abstract thoughts (Margulies et al., 2016). The DMN extreme of the gradient was of particular interest in the present study. Previous studies have linked DMN with general semantic processing (Binder & Desai, 2011; Binder, Desai, Graves, & Conant, 2009; Binder et al., 1999), and it is thought to play an important role in constructing a mental representation of discourse during comprehension and production (often termed a “situation model”, Garrod & Pickering, 2004; Heidlmayr et al., 2020; Kintsch & Van Dijk, 1978).

**Figure 3.**
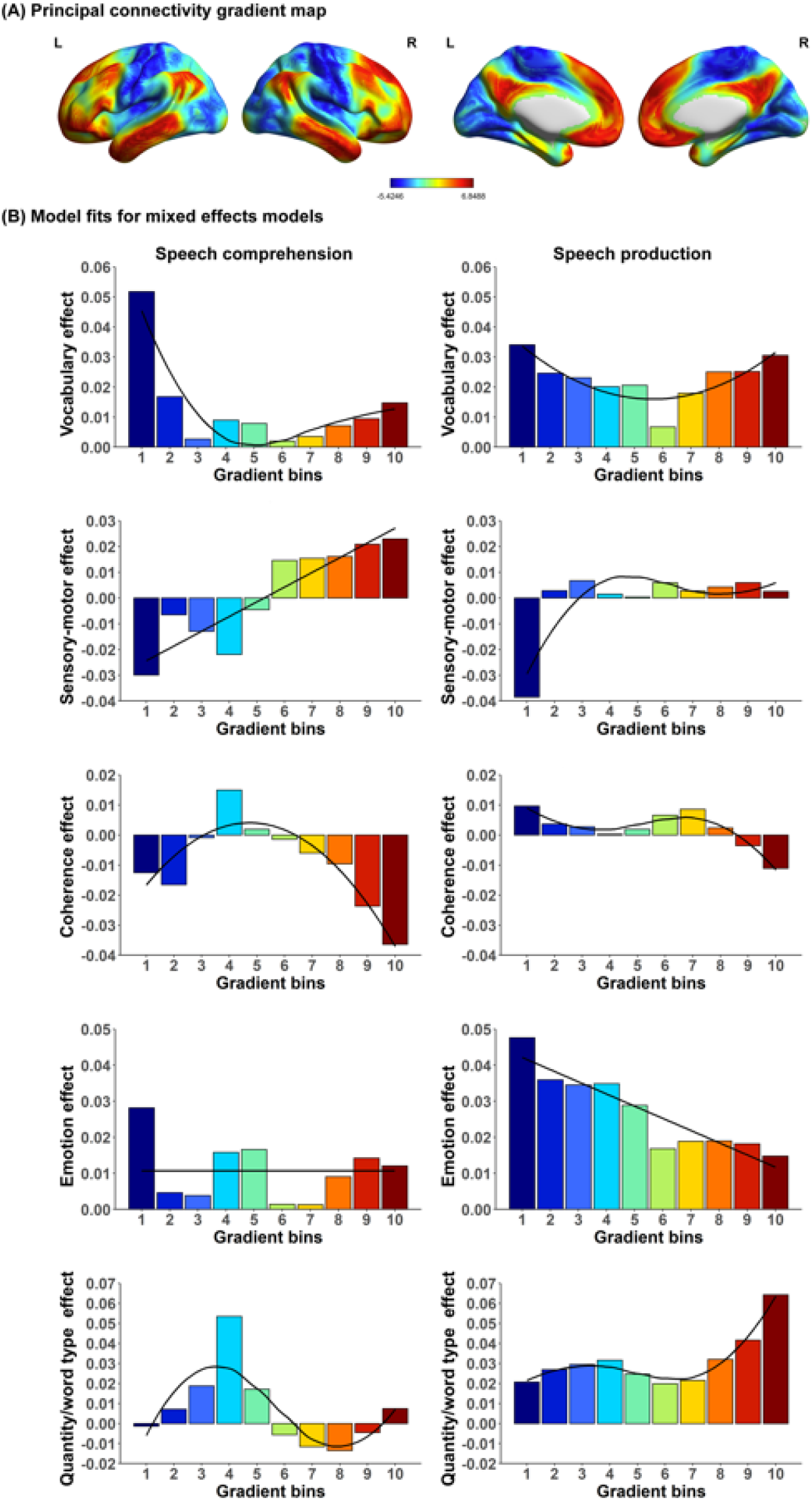
(A) Principal connectivity gradient map from Margulies et al. (2016), with the unimodal extreme of the gradient shown in blue and the DMN extreme in red. (B) Model fits for the mixed effects models that examined the effect of the gradient on psycholinguistic PCs activation during speech comprehension and production. Linear, quadratic and cubic models were fitted for each PC. The trend line for the best-fitting model (lowest AIC) in each case is shown.

We aimed to investigate how the effects of psycholinguistic properties varied along this gradient. To do this, we divided voxels along the gradient into 10 equally-sized bins (from sensorimotor to DMN regions) using similar methods to Xiuyi Wang, Margulies, Smallwood, and Jefferies (2020). Then we extracted mean PC effects (beta values) for each participant for the voxels contained in each bin. To determine how the effects varied along the cortical gradient, we fitted linear mixed models predicting these psycholinguistic effects for comprehension and production separately. Our fixed effect was the position of the bin on the gradient (from 1 to 10), treated as a numeric variable and centered and scaled. To test for higher order relationships between gradient bins and psycholinguistic effects, we also built mixed models which included second-order and third-order terms for bin position. We investigated if the including of higher orders of predictors improved model fits, by comparing the Akaike information criterion (AIC) of different models. We selected the model with the lowest AIC to represent the change in effects along the gradient. All models in this section included random intercepts by participant.

## 3. Results

### 3.1. Characteristics of speech

The relationships among the 13 psycholinguistic properties of interest were first investigated with correlation analysis. We divided each 50 s speech period into ten 5-second blocks and calculated the psycholinguistic properties for each block. Then we combined the measures of all the speech blocks in both comprehension and production tasks and computed Pearson correlations for each pair of properties. Table 1 shows the results of the correlation analysis. As expected, none of the properties was entirely independent of the others: every property covaried with some of the other speech properties. For example, speech blocks with lower word frequency typically contained words acquired later in life and had lower semantic diversity. Blocks that included more concrete words were unsurprisingly more perception- and action-related. These results underscore the need to utilize data reduction techniques to generate latent dimensions as measures of speech characteristics.

**Table 1.** Correlations among psycholinguistic properties of speech.

|  | 1. | 2. | 3. | 4. | 5. | 6. | 7. | 8. | 9. | 10. | 11. | 12. |
| --- | --- | --- | --- | --- | --- | --- | --- | --- | --- | --- | --- | --- |
| 1. Frequency | -- |  |  |  |  |  |  |  |  |  |  |  |
| 2. Concreteness | -0.07 | -- |  |  |  |  |  |  |  |  |  |  |
| 3. AoA | -0.50 | -0.41 | -- |  |  |  |  |  |  |  |  |  |
| 4. Number of phonemes | -0.37 | -0.20 | 0.42 | -- |  |  |  |  |  |  |  |  |
| 5. SemD | 0.66 | -0.34 | -0.28 | -0.13 | -- |  |  |  |  |  |  |  |
| 6. Number of words | 0.10 | 0.02 | -0.08 | -0.14 | 0.08 | -- |  |  |  |  |  |  |
| 7. Proportion closed-class words | -0.05 | 0.02 | 0.02 | -0.01 | -0.01 | 0.28 | -- |  |  |  |  |  |
| 8. Global coherence | -0.13 | 0.16 | -0.02 | -0.08 | -0.25 | -0.08 | -0.12 | -- |  |  |  |  |
| 9. Local coherence | -0.12 | 0.15 | -0.04 | -0.09 | -0.19 | -0.005 | -0.003 | 0.46 | -- |  |  |  |
| 10. Arousal | -0.04 | -0.06 | 0.08 | 0.18 | -0.12 | -0.09 | -0.03 | -0.01 | -0.05 | -- |  |  |
| 11. Valence | 0.01 | 0.06 | -0.23 | -0.02 | -0.17 | -0.05 | 0.03 | 0.06 | 0.05 | 0.27 | -- |  |
| 12. Perception strength | 0.09 | 0.60 | -0.43 | -0.12 | -0.18 | -0.04 | 0.01 | 0.11 | 0.09 | 0.14 | 0.32 | -- |
| 13. Action strength | 0.12 | 0.24 | -0.19 | -0.08 | -0.05 | -0.09 | -0.05 | 0.11 | 0.05 | 0.19 | 0.19 | 0.47 |

Note: N=3240, r values greater than +/- 0.034 are significant at $p < 0.05$ (uncorrected).

To further explore the structure among speech properties and generate latent measures of speech characteristics, a PCA was performed on all properties of all speech blocks, which resulted in the extraction of five PCs (which together explained 68% of the variance). The results are reported in Table 2 and example speech passages that scored low and high on each component are presented in Table 3. For convenience, we labelled each component according to the aspects of speech it appeared to index:

PC 1. Complexity of Vocabulary (blocks containing high frequency, high semantic diversity, short, early-acquired words scored highly on this factor while those containing more complex vocabulary received low scores)
PC 2. Sensory-motor content (blocks containing words high in concreteness and sensory motor information scored highly)
PC 3. Coherence (blocks high in global and local coherence loaded positively on this factor)
PC 4. Emotional content (blocks with words high in arousal and valence scored highly)
PC 5. Quantity/word type (blocks containing a high number of words and high proportion of closed class words loaded positively on this factor)

**Table 2.** Results of principal component analysis of speech properties.

|  | PC 1:<br>Vocabulary | PC 2:<br>Sensory-motor | PC 3:<br>Coherence | PC 4:<br>Emotion | PC 5:<br>Quantity/word<br>type |
| --- | --- | --- | --- | --- | --- |
| Frequency | **0.91** | -0.08 | -0.05 | 0.08 | -0.04 |
| SemD | **0.81** | -0.38 | -0.16 | -0.03 | -0.05 |
| AoA | **-0.62** | -0.50 | -0.01 | -0.01 | -0.03 |
| Number of phonemes | **-0.53** | -0.22 | -0.24 | 0.22 | -0.12 |
| Concreteness | -0.19 | **0.94** | -0.06 | -0.24 | 0.01 |
| Perception strength | 0.01 | **0.84** | -0.10 | 0.22 | -0.03 |
| Action strength | 0.11 | **0.45** | -0.03 | 0.37 | -0.17 |
| Local coherence | -0.02 | -0.12 | **0.87** | 0.05 | 0.06 |
| Global coherence | -0.05 | -0.06 | **0.85** | 0.06 | -0.14 |
| Arousal | -0.09 | -0.12 | -0.03 | **0.80** | -0.01 |
| Valence | 0.07 | 0.10 | 0.15 | **0.72** | 0.16 |
| Proportion closed-class words | -0.13 | -0.002 | -0.10 | 0.13 | **0.82** |
| Number of words | 0.11 | -0.06 | 0.02 | -0.02 | **0.77** |

Note: Table shows loadings in pattern matrix following promax rotation. The order of the properties was organized based on their loadings on factors (each property's strongest loading highlighted in bold). AoA = age of acquisition; SemD = semantic diversity.

**Table 3.**
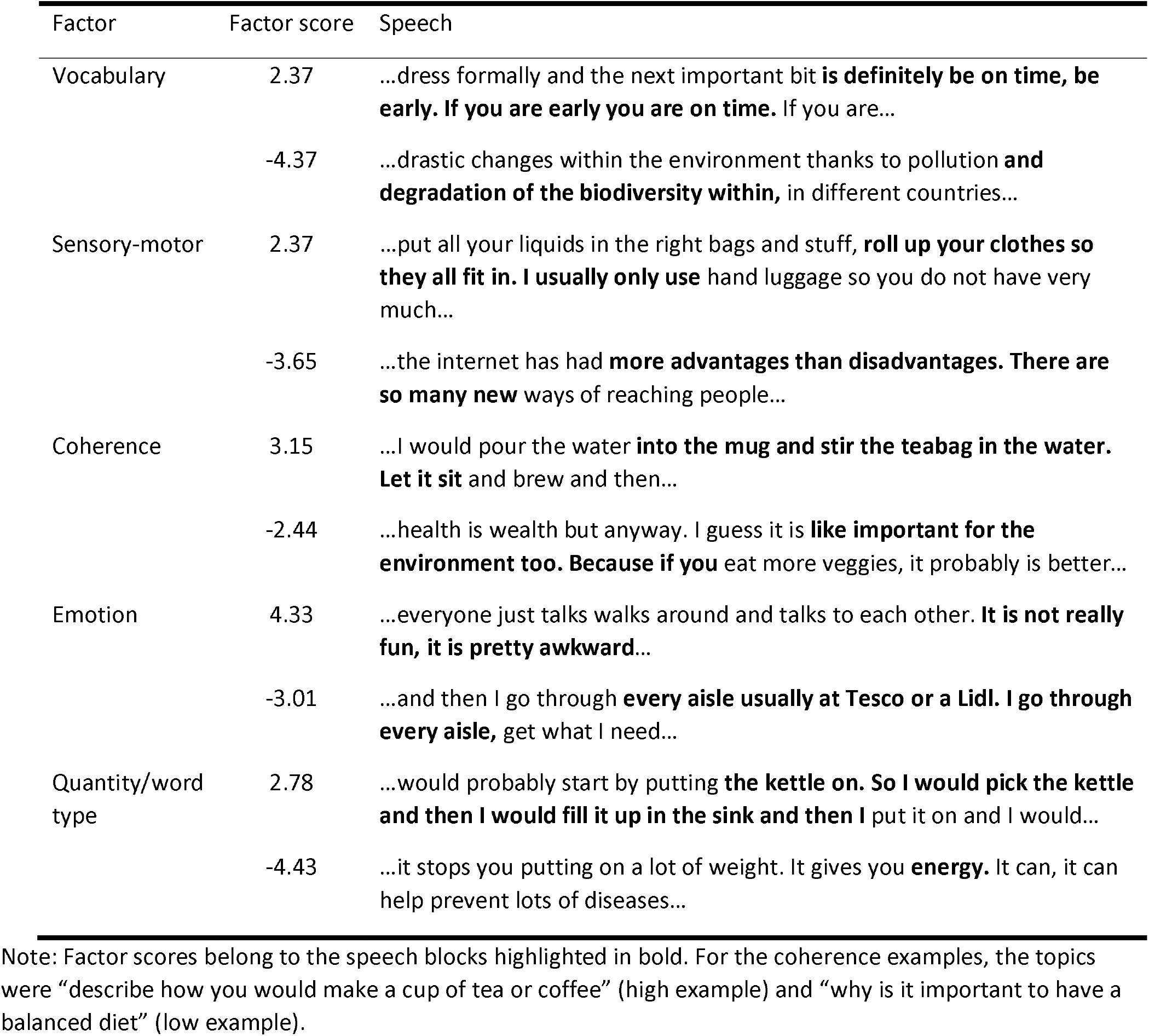
Example speech passages that scored high and low on each psycholinguistic factor.

The PCA indicates that the 13 individual properties we derived from the speech samples cohere into a smaller set of distinct and readily interpretable underlying components. Moreover, these components map clearly onto different levels of language processing: PC1 relates to lexical properties of speech, PC2 and PC4 to semantic content, PC3 to the structure and organisation of the discourse and PC5 to the overall rate of speech and its grammatical content. This result is consistent with previous studies that have used PCA to decompose speech properties in healthy and disordered populations; these have found latent factors relating to similar levels of language processing (Alyahya et al., 2020b; Hoffman, 2019; Hoffman et al., 2018; Mirman et al., 2015). In addition, we conducted two control analyses to investigate the effects of (a) varying the length of the time window over which properties were aggregated, and (b) using lexical properties for all open-class words and not just nouns. These control analyses provided similar results to the main analysis (see Supplementary Tables S2 and S3).

The PCA provides us with an interpretable set of latent properties with which to interrogate our neuroimaging data. Our main neuroimaging analysis was therefore based on the PCA results, in which neural activity during discourse was simultaneously predicted from block scores on the five latent factors described above (as well as the position of the block within the discourse). The results of parallel analyses investigating the effects of the 13 individual language properties are provided in Supplementary Materials for interested readers.

### 3.2. Activation during speech processing as a function of psycholinguistic properties

To investigate how brain activation co-varied with the five PCs underlying speech processing, we performed voxel-wise and network-level analyses. At the voxel level, we computed effect sizes (beta values) for each PC in each voxel across the whole brain, as shown in Figure 2. Activation in hot colour areas was positively correlated with the scores on the five psycholinguistic factors, increasing when participants comprehended or produced speech that scored positively on this factor, whereas cold colour areas were negatively correlated with the factor scores. At the network level, we classified voxels according to their position on the principal cortical gradient reported by Margulies et al. (2016). This gradient places the cortex along a continuum from primary sensory and motor regions to the DMN (see Figure 3A). We computed and plotted the mean effect size of each PC at 10 positions along this gradient, as shown in Figure 3B, along with the model fits based on AIC of the corresponding mixed effects models (Table 4). Taken together, the whole-brain maps and gradient plots provide information about how activation in different brain regions changes as a function of the properties of discourse during both comprehension and production. Finally, to assess the level of convergence between language tasks, we also computed the voxel-wise correlation between the effect size (beta) maps for comprehension and production, as reported in Figure 2 (using similar methods with Morales et al., in press; Xiuyi Wang et al., 2020). In the following section, we provide a verbal summary of the topographic distribution of the effects observed for each PC, focusing particularly on areas known to be involved in language processing and semantic cognition.

**Table 4.**
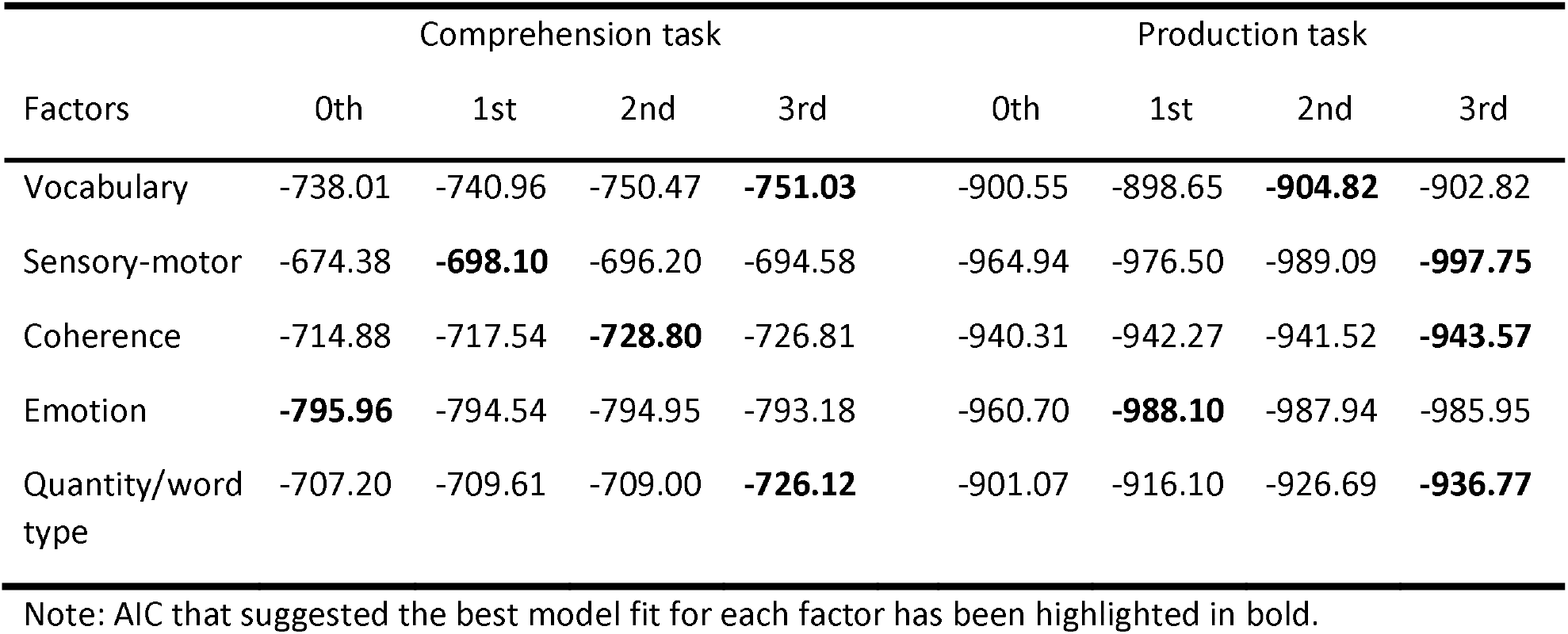
AIC for the mixed effects models with different order effects of the gradient bins on psycholinguistic PCs activation during speech.

| Factors | Comprehension task |  |  |  | Production task |  |  |  |
| --- | --- | --- | --- | --- | --- | --- | --- | --- |
|  | 0th | 1st | 2nd | 3rd | 0th | 1st | 2nd | 3rd |
| Vocabulary | -738.01 | -740.96 | -750.47 | **-751.03** | -900.55 | -898.65 | **-904.82** | -902.82 |
| Sensory-motor | -674.38 | **-698.10** | -696.20 | -694.58 | -964.94 | -976.50 | -989.09 | **-997.75** |
| Coherence | -714.88 | -717.54 | **-728.80** | -726.81 | -940.31 | -942.27 | -941.52 | **-943.57** |
| Emotion | **-795.96** | -794.54 | -794.95 | -793.18 | -960.70 | **-988.10** | -987.94 | -985.95 |
| Quantity/word type | -707.20 | -709.61 | -709.00 | **-726.12** | -901.07 | -916.10 | -926.69 | **-936.77** |

Note: AIC that suggested the best model fit for each factor has been highlighted in bold.

For the vocabulary component, the mean net effect across the cortical gradient was positive in both comprehension and production tasks. That is, there was a tendency towards greater activation for passages that contained less complex nouns. During speech comprehension, increased activation for less complex vocabulary was evident within angular gyrus, superior frontal gyrus, posterior cingulate gyrus, precuneus, and areas in the occipital lobe. This interpretation was supported by the gradientbased analysis, which showed a strong positive effect at the unimodal end of the gradient (corresponding to visual cortex in the occipital lobe), and an increase in positive effects towards the opposing DMN end of the spectrum. In contrast, when participants heard more complex terms, language areas showing activation increases included inferior frontal gyrus and inferior temporal gyrus and the ventral anterior temporal lobe. Similar sets of regions showed effects of the vocabulary component in the speech production task. Indeed, we found a positive correlation between the activation maps of the vocabulary component in comprehension and production tasks (r = 0.25) suggesting convergence across language tasks.

For the sensory-motor factor, during speech comprehension stronger activation for more concrete content included parts of the DMN such as the angular gyrus, posterior cingulate, and ventromedial prefrontal cortex, as well as parahippocampal gyrus, inferior temporal gyrus and inferior frontal gyrus. In contrast, there was strong activation increase for abstract words in areas of superior temporal gyrus close to and anterior to primary auditory cortex, as well as in occipital cortex. In support of this interpretation, the gradient analysis for comprehension showed a linear trend, with cortex closer to primary sensory-motor regions activating to more abstract content while regions towards the DMN activating to more concrete content. During speech production, the effect for concrete words in DMN appeared less pronounced (in line with no consistent effect in the gradient analysis) and there was a pronounced increase for more abstract content in the left anterior temporal lobe. Despite some apparent differences, the correlation between comprehension and production for the sensory-motor factor was the strongest of all five PCs (r = 0.42), suggesting that the sensory-motor content of discourse has relatively similar neural correlates whether one is speaking or listening.

For the coherence factor, superior temporal gyrus and parts of the left frontal lobe showed greater activation when people heard more coherent passages of speech. However, stronger correlations were observed for less coherent speech, in DMN regions such as the angular gyrus, posterior cingulate and ventromedial prefrontal cortex, and along the length of the superior temporal sulcus. Similarly, the gradient-based analysis indicated that the strongest negative response to coherence was at the DMN extreme of the cortical gradient. For the speech production task, the gradient analysis showed weaker effects and the negative effects of coherence were much less pronounced. There were, however, notable positive effects in inferior frontal gyrus, posterior middle temporal gyrus and the inferior parietal lobule, all of which showed higher activation for more coherent speech. Less coherent discourse was also associated with greater activation in DMN regions during production, though to a lesser extent than in the comprehension task. The activation maps in comprehension and production were positively correlated with each other (r = 0.29), suggesting some convergence in the distribution of effects. These results are similar to our previous analysis of coherence in this dataset (Morales et al., in press).

For the emotion component, the gradient analysis revealed net positive effects of emotional content on brain activation, particularly during the production task. Areas showing activation increases during comprehension of high-emotion passages included cingulate gyrus, insula, medial frontal gyrus, precuneus and superior temporal gyrus. In contrast, inferior frontal gyrus and parahippocampal gyrus were more activated when speech included more neutral content. For the production task, parts of the cingulate gyrus, insula, medial frontal gyrus, superior temporal gyrus and precentral/postcentral gyri showed greater activation to high-emotion speech. Inferior parietal lobule, angular gyrus, inferior frontal gyrus and parts of the ventral temporal lobe showed greater activation to neutral speech, though these effects were small. A positive correlation was found when the effect maps of the emotion factor in the two tasks were compared but this was very weak (r = 0.08), suggesting that the processing of emotional language engages somewhat different systems depending on whether one is speaking or listening to speech.

Finally, for the quantity/word type factor, in the comprehension task, a robust increase to speech including more (closed-class) words was observed in superior temporal gyrus and superior temporal sulcus bilaterally, which was the strongest effect for this component. The gradient analysis for comprehension showed a pronounced positive effect around gradient bin 4, which corresponds to the position of mid-to-anterior STG regions on the gradient. Greater activation to blocks including fewer words and fewer closed-class words was found in intraparietal sulcus, inferior frontal gyrus and insula, with larger effects in the right hemisphere. These effects may indicate regions that deactivate during language comprehension. In the production task, gradient analysis indicated a net positive effect on activation across the brain when participants produced more words, with the strongest effect towards the DMN. In the effect maps, particularly strong effects were seen in precuneus, superior frontal gyrus, angular gyrus, the lateral temporal lobe and motor cortices. No brain regions activated more strongly for speech blocks containing fewer (closed-class) words, with the possible exception of the ventral anterior temporal cortices. When comparing the activation maps in comprehension and production tasks, we found a weak positive correlation for the quantity/word type factor (r = 0.18).

We also generated topographic activation maps for each of the individual psycholinguistic properties, to explore raw feature effects without accounting for their relationships with other psycholinguistic properties. These results are provided in Supplementary Materials.

## 4. Discussion

A common approach in language neuroscience is to investigate how neural activity is influenced by the properties of the language being processed. Most fMRI studies to use this method have used simple experimental stimuli at the single word or sentence level, rather than more naturalistic discourse. In addition, few studies have investigated psycholinguistic effects during language production rather than comprehension. In this exploratory fMRI study, we investigated how neural activity covaried with the psycholinguistic properties of naturalistic discourse, comparing speech comprehension with production in the same participants. Several findings emerged from the present study. First, we found that PCA could be used successfully to reduce a broad range of linguistic properties of interest to five meaningful latent factors, which quantified the lexical properties of speech (vocabulary), its semantic content (sensory-motor, emotion), its organisation at the discourse level (coherence) and its overall quantity and composition. Second, by exploring the neural correlates of each of these psycholinguistic factors at a whole-brain level, we found frequent convergence between effects observed in language comprehension and production, though there was also evidence for divergence on some properties. Third, we observed different responses across the brain to different factors, some of which corroborate previous findings derived from more constrained experimental stimuli, and some of which suggest new hypotheses for future research. Overall, our study demonstrates that naturalistic fMRI paradigms can be used to study neural processes in speech production as well as comprehension. In this Discussion, we note where our findings are most compatible with the existing literature and where they generate new research questions and hypotheses for future research.

We begin by considering the level of convergence between effects in comprehension and production. The strongest correlation between tasks was for the sensory-motor speech factor. This factor indexes the degree to which speech passages contain concrete referents associated with perception and action and therefore loads most squarely on the semantic level of language processing. Most theories of language agree that semantic representations are shared between comprehension and production (Gambi & Pickering, 2017; Garrod & Pickering, 2004; Hagoort, 2013; Hickok & Poeppel, 2007; Kintsch & Van Dijk, 1978; Levelt et al., 1999; Pickering & Garrod, 2021). In line with this general consensus, the sensory-motor component showed the highest degree of convergence between our two tasks, suggesting that sensory-motor content of speech influences neural activity in a similar fashion whether one is speaking or listening to speech. In contrast, the lowest correlation between tasks occurred for the emotion factor. This result suggests that there may be important differences in the neural systems engaged by emotional content in one’s own language production, when compared with someone else’s speech. Studies investigating theory of mind have found neural differences when making judgements about our own emotional states compared with inferring and empathising with the mental states of other people (Ochsner et al., 2004; Reniers, Völlm, Elliott, & Corcoran, 2014; Ruby & Decety, 2004). Here we have found that, even during passages of relatively neutral discourse, the brain’s response to emotion words seems to vary depending on their source. Future studies could investigate the precise neural loci of such effects and the underlying mechanisms that give rise to these differences. It is possible, for example, that we experience an increased propensity to simulate or re-experience emotional states when we recount our own experiences, compared with listening to other people’s. The correlation for the quantity factor was also low. This is perhaps unsurprising since variations in speech rate are likely to place demands on different sensorimotor systems depending on whether one is listening to or producing speech. The remaining two factors (vocabulary and coherence) showed intermediate levels of correlation between the two tasks, which suggests some overlap between comprehension and production.

In addition to measuring whole-brain convergence between comprehension and production, the topographic distribution of different psycholinguistic effects can provide useful insights into the engagement of neural systems during discourse processing. For the vocabulary factor, consistent with the positive effects of word frequency in previous studies of single-word reading (Carreiras, Riba, Vergara, Heldmann, & Münte, 2009; Graves et al., 2010; Prabhakaran, Blumstein, Myers, Hutchison, & Britton, 2006), we found that simpler vocabulary was associated with increased activation in DMN areas including angular gyrus, cingulate gyrus and precuneus, which have been strongly implicated in semantic processes (Binder et al., 2009). One can intuit that simpler vocabulary is more likely to elicit stronger activation in a semantic network due to their extensive exposure compared with more complex words, which are lower in frequency, acquired later in life and have fewer links to other words (Binder & Desai, 2011; De Deyne & Storms, 2008; Reilly & Desai, 2017). In contrast, the negative correlations for word frequency observed previously in inferior frontal gyrus and ventral temporal cortex during single-word reading tasks (Carreiras et al., 2009; Graves et al., 2010; Hauk et al., 2008; Hoffman, Lambon Ralph, & Woollams, 2015; Prabhakaran et al., 2006) were also observed in our study when participants processed more complex spoken language. Given the established role of inferior prefrontal cortex in cognitive control, particularly during language tasks (Jackson, 2021), it is not surprising that the processing of complex language engages this area. The greater engagement of ventral anterior temporal regions to more complex vocabulary was less expected and warrants future investigation. Both the ventral anterior temporal cortex and inferior parietal cortex have been proposed as the sites of semantic “hubs” that code conceptual knowledge (Binder & Desai, 2011; Lambon Ralph, Jefferies, Patterson, & Rogers, 2017; Mirman, Landrigan, & Britt, 2017). Here, however, they showed opposite responses to the vocabulary component of discourse. Further investigation of the specific factors that influence discourse-related activity in each region may be valuable in teasing apart the specific functions of these regions.

Our second factor indexed the sensory-motor content of language. This factor has been frequently studied by manipulating the concreteness of written words. Such studies have reliably found that angular gyrus, precuneus/cingulate, parahippocampal gyrus and ventromedial prefrontal cortex (generally considered DMN regions) show increased activation to more concrete words (for meta-analyses, see Bucur & Papagno, 2021; J. Wang et al., 2010). Our data largely replicate these effects in auditory comprehension, though they appeared somewhat weaker during language production. Concrete concepts are thought to have a richer and more easily accessed semantic representation (Paivio, 1991; Schwanenflugel, 2013), thus the activation of the above DMN areas could reflect engagement of richer semantic representations for speech containing more concrete content (Binder & Desai, 2011). Our results are also consistent with the findings that emerge from the studies focusing on examining effects of sensory-motor information on brain activity. For example, Fernandino et al. (2016) found that multiple sensory-motor attributes of words can modulate the neural activity in a similar set of regions to our study, including parahippocampal gyrus, precuneus/cingulate, medial prefrontal cortex and angular gyrus. Other studies have suggested that left lateral occipitotemporal cortex is critically involved in representing action information (Wu, Wang, Wei, He, & Bi, 2020; Wurm & Caramazza, 2019; Wurm, Caramazza, & Lingnau, 2017), which was one of the properties that contributed to our sensory-motor factor. Consistent with this account, we observed a positive response to more sensory-motor language content in posterior occipitotemporal regions during both language tasks.

Previous meta-analyses have identified greater activation to more abstract word comprehension in left lateral temporal (particularly anterior) regions and the left IFG (Bucur & Papagno, 2021; J. Wang et al., 2010). Consistent with this, we observed greater activation to less sensory-motor speech passages in the lateral temporal cortices in both comprehension and production. However, there was no suggestion of a similar effect in IFG for either task. We suggest that this is potentially an important point of divergence between experimental stimuli and more natural speech, which warrants future investigation. It has been proposed that abstract concepts engage IFG because they have greater contextual variability than concrete concepts (Schwanenflugel & Shoben, 1983) and thus require more engagement of semantic selection and control processes supported by IFG (Hoffman, Binney, et al., 2015; Hoffman, Jefferies, & Lambon Ralph, 2010; Noppeney & Price, 2004). In the present paradigm, unlike in previous studies, abstract words were embedded in naturalistic discourse and their interpretation was therefore constrained by the rich prior context of the discourse. Therefore, it is possible that the executive demands for abstract words in our study were minimised, which could account for the lack of IFG response to this variable. This interpretation supports the context availability theory of concreteness effect (Schwanenflugel & Shoben, 1983) and suggests a need to directly compare the semantic processing mechanisms of discourse versus single words in future studies. We suggest that the executive control demands for different types of concepts could vary radically according to the specific experimental environment in which they are presented.

The coherence factor in our data loaded on measures of global coherence, which indexes the degree to which utterances conform to the expected topic of the discourse, and local coherence, which measures the relatedness of neighbouring passages of speech (Glosser & Deser, 1992). Both of these measures reflect the high-level organisation of discourse. The results for this factor replicate and extend our previous analyses of the current dataset, in which we investigated the effect of global coherence specifically (Morales et al., in press). Our previous study found that DMN regions showed greater activation when less coherent speech was heard or produced, potentially reflecting updating of mental representations when discourse deviated from the expected topic. The present results support this conclusion and demonstrate that the previous findings are valid even when the influences of other properties of language are statistically controlled with PCA.

For the emotion factor, several areas showed increased activation to speech containing more emotional content, including anterior cingulate cortex, insula, and medial frontal gyrus. These areas were also reported in previous studies as typically responding to emotionally significant language or more general stimuli (e.g., pictures) (Citron, 2012; Kensinger & Schacter, 2006; Vigliocco et al., 2014). Nevertheless, another classical emotion-related area, the amygdala, showed a slight negative effect to emotional language in the present study. As stated earlier, effects of the emotion factor were very weakly correlated across comprehension and production, suggesting that the neural processing of emotional discourse during language production differs in important ways from in language comprehension. In general, brain regions were much more likely to show emotion-related activation increases during the production task. The reasons for this are unclear. One possibility is that our comprehension task did not elicit a high level of emotional engagement from participants, since they did not see the face of the speaker and was not aware of their identity. It is important to also note that our topics were designed to probe general knowledge rather than personal experiences, and thus were not suited to eliciting highly emotional discourse. Thus our data may be less sensitive to this aspect of language. In other words, the variance of neural activity elicited by the emotion factor could be harder to detect than the other aspects of language, as there was not much emotional content in the stimuli.

Lastly, for the quantity/word type factor, strong activation increases were observed in auditory cortices and the surrounding superior temporal gyri during comprehension. Passages loading on this factor contained a high number of words and a high proportion of closed-class words that carry syntactic rather than semantic information. This result is consistent with other studies that have investigated effects of speech rate on brain activation (Dhankhar et al., 1997; Mummery et al., 1999; Price et al., 1992) and likely reflects the fact that regions surrounding primary auditory cortex play a critical role in speech perception (Hickok & Poeppel, 2007). The superior temporal gyrus is also implicated in syntactic processing (Friederici, 2012), which may also explain why this region responds strongly to passages containing a large number of closed-class words. Strong negative effects of quantity/word type during comprehension were observed in large swathes of the right hemisphere, which may indicate increasing deactivation of brain areas not responsive to language, with deactivation occurring in proportion to the intensity of speech processing demands.

To our knowledge, no neuroimaging study has previously investigated the parametric effect of speech quantity in discourse production. Effects here were somewhat different to those observed in comprehension, with increasing activation as a function of speech quantity/word type across much more of the cortex. Activation increases were strongest in the angular gyri bilaterally, with other DMN regions also showing strong effects. As we have noted earlier, these regions are implicated in semantic representation, mental models of events and situations and with self-generated thought more generally (Andrews-Hanna, Smallwood, & Spreng, 2014; Binder & Desai, 2011; Margulies et al., 2016). One hypothesis arising from this result is that the more strongly people engage these systems, the more fluently and rapidly they are able to generate discourse.

## 5. Conclusions

In conclusion, using PCA to derive measures of language properties, the current study is one of the first to directly compare the neural correlates of psycholinguistic effects in naturalistic discourse during comprehension and production. Findings of this exploratory study suggest a number of directions for future work. First, previous work has not explicitly investigated to what extent the neural correlates of psycholinguistic properties overlap during speech comprehension and production. Our results suggest that the alignment of discourse processes during listening and speaking is complex, since their neural correlates were similar across listening and speaking for some aspects of speech (e.g., the sensory-motor factor) but not others (e.g., the emotion factor). Future studies should investigate why different aspects of speech elicit different degrees of neural alignment between comprehension and production.

Second, despite the demonstrable benefits of naturalistic paradigms (Hamilton & Huth, 2020; Hasson & Honey, 2012; Nastase et al., 2020; Yarkoni et al., 2008), most fMRI studies still rely on relatively non-naturalistic single-word designs, with small sets of stimuli. Our results suggest that some findings obtained with experimentally constrained paradigms may not be generalized to more naturalistic language processing. Therefore, ecological validity needs to be taken into account in future studies, and the underlying mechanisms leading to differences between naturalistic and non-naturalistic language processing should be investigated. Of course, the “naturalness” of a stimulus is a matter of degree. We used discourse passages of a limited 50 s duration, where participants were instructed to discuss particular topics. This could be argued to be somewhat artificial. For comprehension, open fMRI datasets using more much longer stimuli like narrated stories are available (Li et al., 2021; Nastase et al., 2021). We are not aware of similar data for discourse production and we believe the field would benefit greatly from such data.

Finally, in the present study, the structure of psycholinguistic properties was explored using PCA, which is a data-driven decomposition technique that has been previously used with success in structural MRI studies and fMRI studies (Alyahya et al., 2020a, 2020b; Fernandino et al., 2016; Hauk et al., 2008; Huth et al., 2016; Mirman et al., 2015). Here, we successfully implemented this technique with naturalistic discourse data during both comprehension and production to generate predictors for fMRI data, which provided strong evidence for the applicability of this technique in future studies. Thus, our findings not only contribute to understanding of shared and distinct neural processes in the comprehension and production of naturalistic discourse, but also provide evidence for the utility of applying quantitative analysis of naturalistic speech to study the neural mechanisms of language.

## Open Practices

Data for this study are available at https://osf.io/4tbpm/. Activation maps for each psycholinguistic component and single psycholinguistic property are provided at individual level and group level in the data repository. The conditions of our ethics approval do not permit public archiving of the raw MRI images associated with this study or the speech responses, which include ethically sensitive content like personal experiences of the participants. Readers seeking access to this data should contact the PI Dr Paul Hoffman, or the local ethics committee at the University of Edinburgh. Access will be granted to named individuals in accordance with ethical procedures governing the reuse of sensitive data. Specifically, the following conditions must be met to obtain access to the data: approval by the Psychology Research Ethics Committee of the University of Edinburgh and a suitable legal basis for the release of the data under GDPR. No part of the study procedures or analysis plans was preregistered prior to the research being conducted.

## CRediT authorship contribution statement

Wei Wu: Conceptualization, Formal analysis, Validation, Writing - original draft, Writing - review & editing, Visualization. Matías Morales: Investigation, Formal analysis, Writing - review & editing. Tanvi Patel: Investigation, Formal analysis, Writing - review & editing. Martin J. Pickering: Conceptualization, Writing - review & editing. Paul Hoffman: Conceptualization, Writing - review & editing, Project administration, Supervision, Funding acquisition.

## Acknowledgements

The project was supported by a BBSRC grant (BB/T004444/1). Imaging was carried out at the Edinburgh Imaging Facility (www.ed.ac.uk/edinburgh-imaging), University of Edinburgh, which is part of the SINAPSE collaboration (www.sinapse.ac.uk). We are grateful to the University of Minnesota Center for Magnetic Resonance Research for sharing their neuroimaging sequences.

## Declarations of interest

none.

## Notes

### Competing Interest Statement

The authors have declared no competing interest.

### Summary of Updates

We only made two significant changes in this version: 1. Added an extra sentence in Figure 3B legend (i.e., "Linear, quadratic and cubic models were fitted for each PC. The trend line for the best-fitting model (lowest AIC) in each case is shown."). 2. Deleted one statement in the Open Science section, as the editor suggested that it was not expected and not required for Exploratory Reports (i.e., "We report how we determined our sample size, all data exclusions, all inclusion/exclusion criteria, whether inclusion/exclusion criteria were established prior to data analysis, all manipulations, and all measures in the study").

