## Supplementary Materials for "Modulation of brain activity by psycholinguistic information during naturalistic speech comprehension and production"

**Supplementary Information**

**Overview**

| Supplementary Figure S1 | Distribution of the PC scores of the psycholinguistic factors. |
| --- | --- |
| Supplementary Figure S2 | Unthresholded maps of single psycholinguistic property effects (beta values) on speech comprehension and speech production. |
| Supplementary Figure S3 | Model fits for the mixed effects models that examined the effect of the gradient on psycholinguistic properties activation during speech comprehension and production. |
| Supplementary Table S1 | Prompts for speech comprehension and production. |
| Supplementary Table S2 | Results of principal component analysis of speech properties based on 10-second speech blocks. |
| Supplementary Table S3 | Results of principal component analysis of speech properties based on all open-class words. |
| Supplementary Table S4 | AIC for the mixed effects models with different order effects of the gradient bins on psycholinguistic properties activation during speech. |

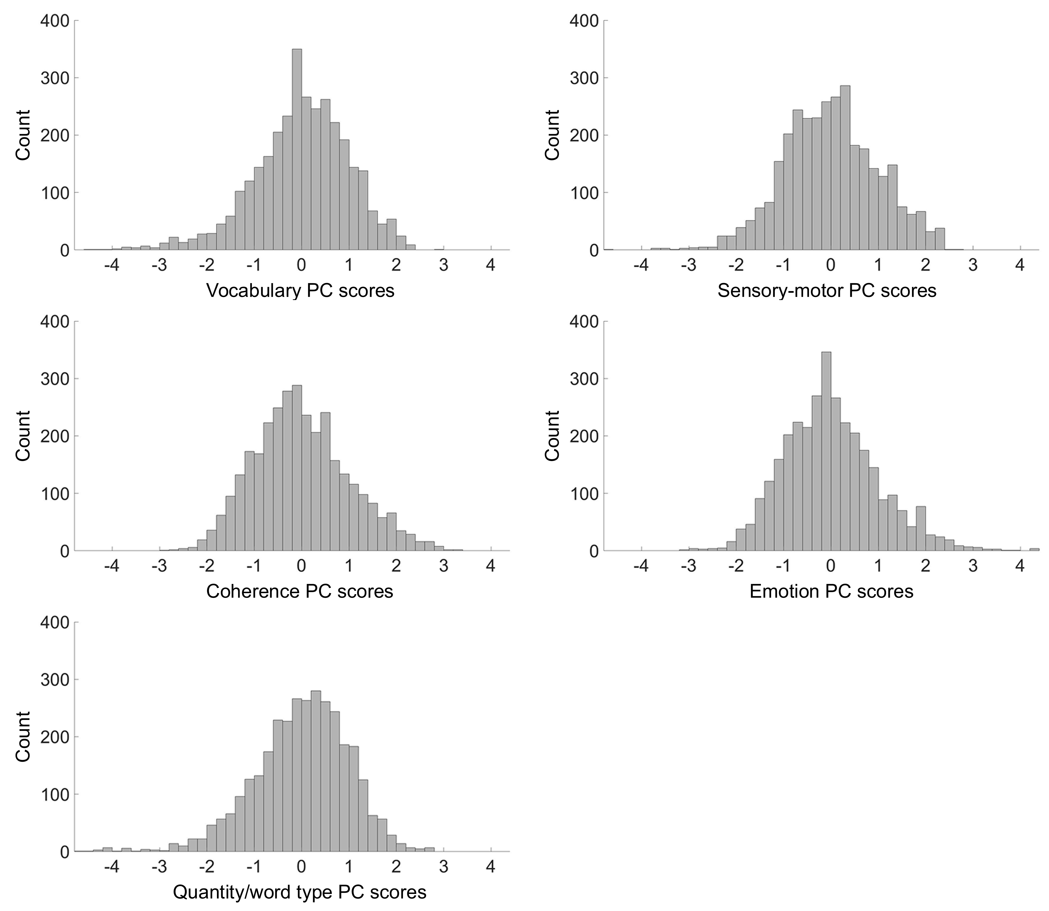

**Figure S1. Distribution of the PC scores of the psycholinguistic factors. The height of the bars indicates the count of the 5-second speech blocks whose PC scores fell within a specified range.**

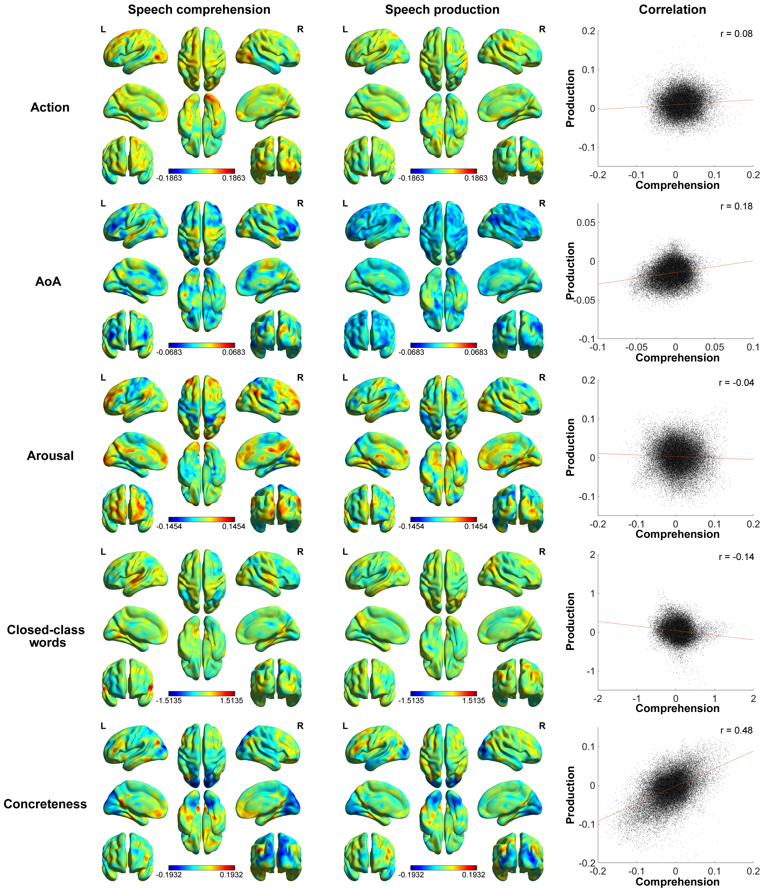

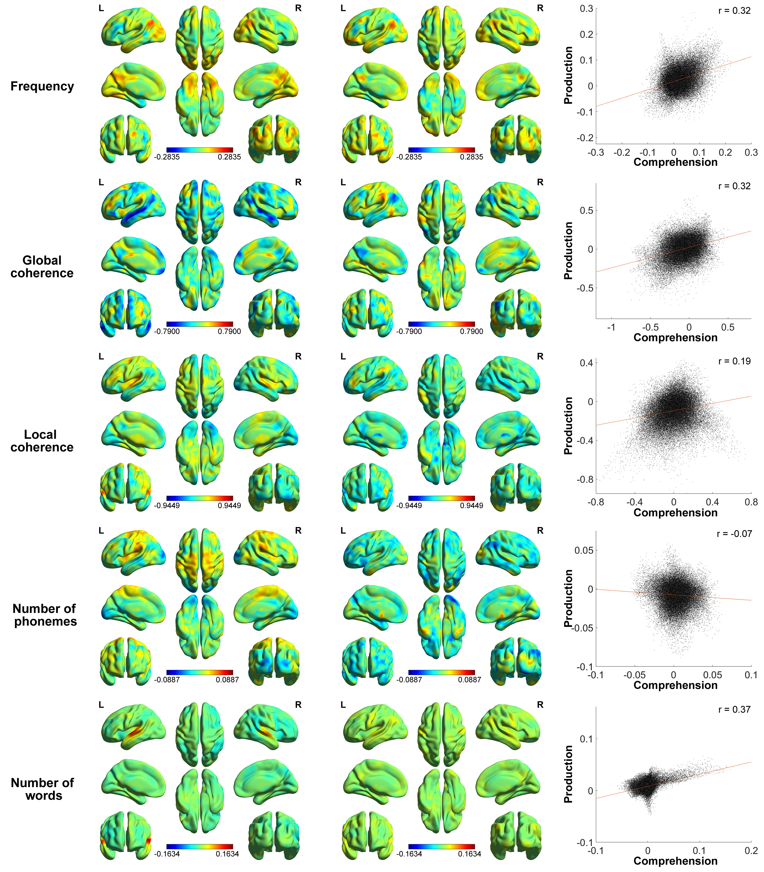

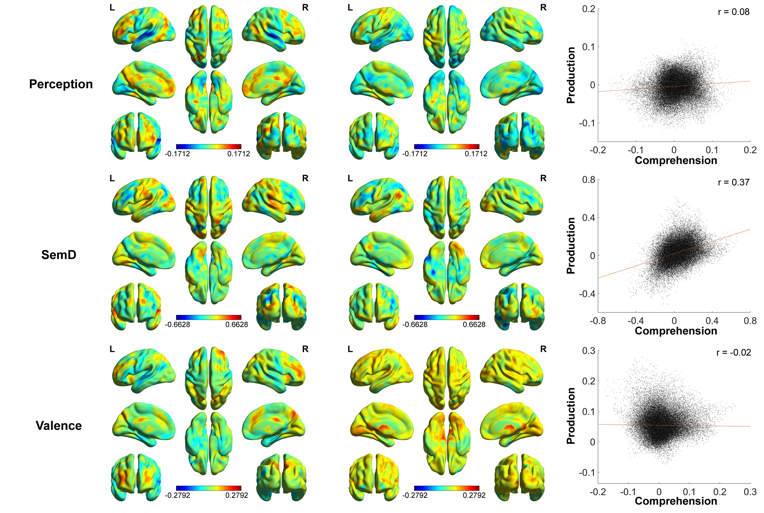

**Figure S2. Unthresholded maps of single psycholinguistic property effects (beta values) on speech comprehension and speech production.**

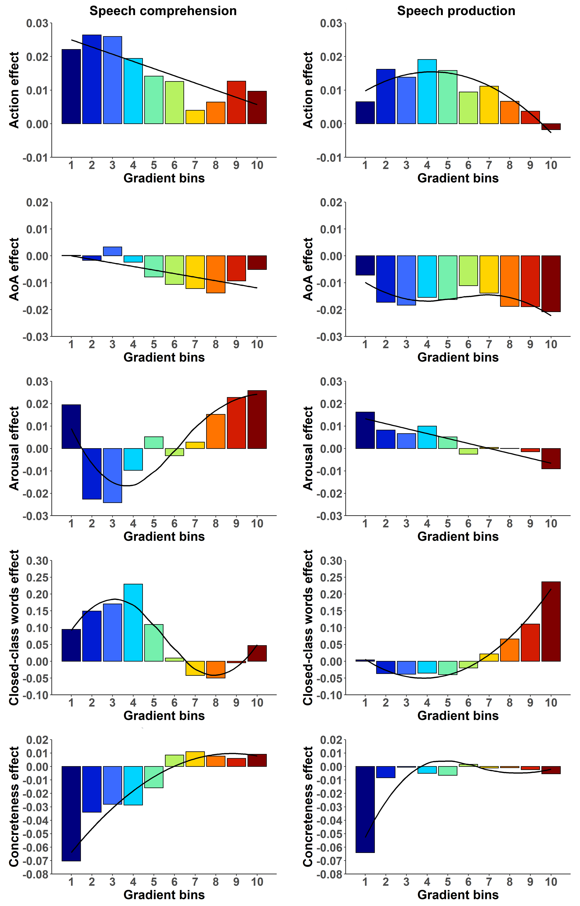

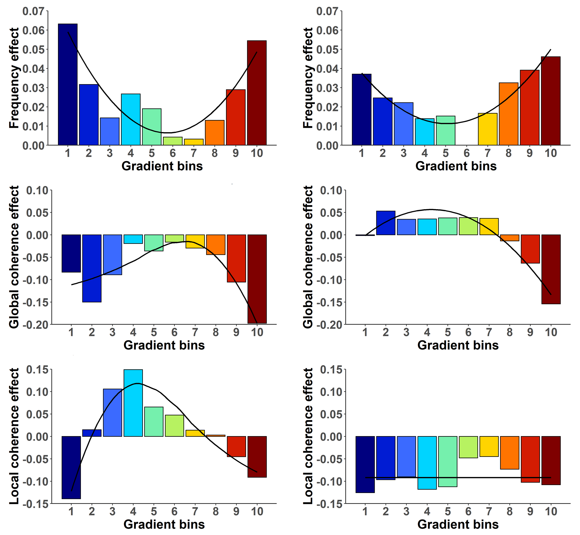

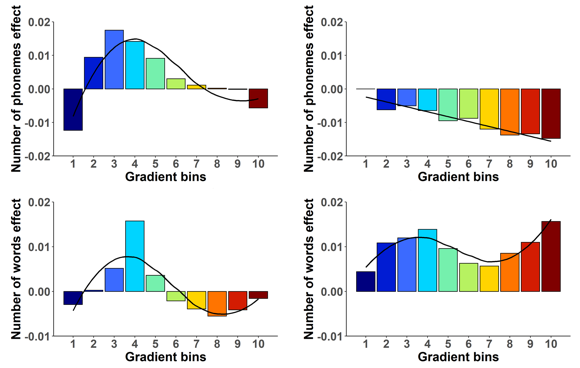

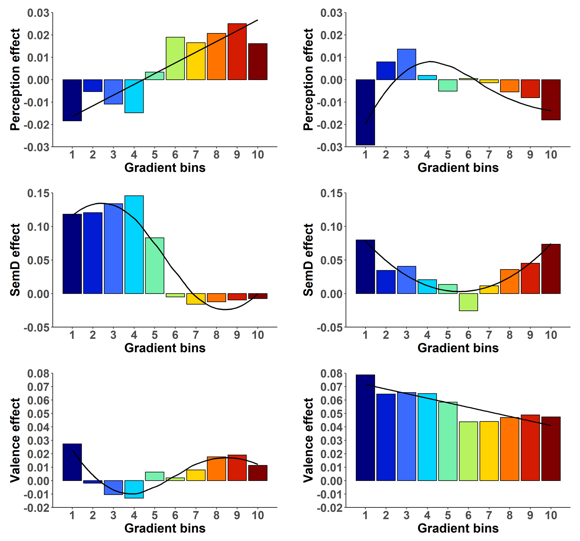

**Figure S3. Model fits for the mixed effects models that examined the effect of the gradient on psycholinguistic properties activation during speech comprehension and production.**

**Table S1. Prompts for speech comprehension and production.**

| Number | Prompts |
| --- | --- |
| Speech comprehension | |
| 1 | What would it be like to live in Antarctica? |
| 2 | What happens when a storm is forecast in the UK? |
| 3 | What do the police do when a crime has been committed? |
| 4 | Which is your favorite season and why? |
| 5 | What do you like or dislike about Christmas? |
| 6 | Do you think it's a good idea to send people to live on Mars? |
| 7 | Why do people come to Scotland on holiday? |
| 8 | What sort of things do you have to do to look after a dog? |
| 9 | Describe a typical visit to a restaurant. |
| 10 | What are the advantages and disadvantages of going to university? |
| 11 | What do people usually do when getting ready for work in the morning? |
| 12 | Describe the steps you would need to take if going somewhere by train. |
| Speech production | |
| 1 | What do people usually do on New Year’s Eve in the UK? |
| 2 | What would you recommend doing during a job interview? |
| 3 | What sorts of things do people do to cope with stress? |
| 4 | Why is it important to have a balanced diet? |
| 5 | Describe the steps you’d take to order-in food. |
| 6 | Describe a typical visit to a grocery store. |
| 7 | What sort of things usually happen at a wedding? |
| 8 | How would you prepare to go on holiday? |
| 9 | Why are some people concerned about climate change? |
| 10 | Do you think the internet has improved people's lives? |
| 11 | What sort of things does a teacher do when at work? |
| 12 | Describe how you would make a cup of tea or coffee. |

**Table S2. Results of principal component analysis of speech properties based on 10-second speech blocks.**

|  | PC 1: Sensory-motor | PC 2: Vocabulary | PC 3: Coherence | PC 4: Emotion | PC 5: Quantity/word type |
| --- | --- | --- | --- | --- | --- |
| Concreteness | **0.91** | -0.31 | -0.06 | -0.20 | 0.0004 |
| Perception strength | **0.87** | -0.08 | -0.10 | 0.21 | -0.05 |
| AoA | **-0.65** | -0.48 | 0.02 | 0.02 | -0.03 |
| Action strength | **0.43** | 0.11 | 0.04 | 0.37 | -0.23 |
| Frequency | 0.005 | **0.93** | -0.03 | 0.10 | -0.03 |
| SemD | -0.35 | **0.84** | -0.14 | 0.001 | -0.02 |
| Number of phonemes | -0.37 | **-0.44** | -0.20 | 0.29 | -0.08 |
| Local coherence | -0.11 | -0.04 | **0.93** | 0.04 | 0.08 |
| Global coherence | -0.05 | -0.09 | **0.90** | 0.05 | -0.12 |
| Arousal | -0.14 | 0.02 | 0.03 | **0.83** | -0.01 |
| Valence | 0.32 | 0.06 | 0.08 | **0.60** | 0.21 |
| Proportion closed-class words | -0.01 | -0.16 | -0.10 | 0.13 | **0.80** |
| Number of words | -0.06 | 0.12 | 0.07 | -0.04 | **0.77** |

Note: Table shows loadings in pattern matrix following promax rotation. The order of the properties was organized based on their loadings on factors (each property’s strongest loading highlighted in bold). AoA = age of acquisition; SemD = semantic diversity. The five PCs together explained 70% of the variance within the set of properties.

**Table S3. Results of principal component analysis of speech properties based on all open-class words.**

|  | PC 1: Sensory-motor | PC 2: Vocabulary | PC 3: Coherence | PC 4: Quantity/word type | PC 5: Emotion |
| --- | --- | --- | --- | --- | --- |
| Concreteness | **0.95** | -0.13 | 0.01 | 0.01 | -0.16 |
| Perception strength | **0.88** | -0.13 | -0.06 | -0.07 | 0.14 |
| SemD | **-0.67** | -0.35 | -0.15 | -0.05 | 0.06 |
| Action strength | **0.56** | -0.19 | -0.13 | 0.07 | 0.34 |
| AoA | -0.30 | **0.90** | -0.01 | 0.05 | -0.03 |
| Number of phonemes | -0.06 | **0.83** | -0.07 | -0.09 | 0.20 |
| Frequency | -0.43 | **-0.68** | -0.05 | 0.01 | 0.15 |
| Local coherence | -0.01 | -0.04 | **0.86** | 0.10 | 0.03 |
| Global coherence | -0.01 | -0.02 | **0.84** | -0.09 | 0.09 |
| Proportion closed-class words | 0.06 | 0.09 | -0.03 | **0.88** | 0.08 |
| Number of words | -0.10 | -0.25 | 0.06 | **0.65** | -0.08 |
| Arousal | -0.01 | 0.29 | -0.02 | 0.10 | **0.80** |
| Valence | -0.003 | -0.17 | 0.18 | -0.09 | **0.63** |

Note: Table shows loadings in pattern matrix following promax rotation. The order of the properties was organized based on their loadings on factors (each property’s strongest loading highlighted in bold). AoA = age of acquisition; SemD = semantic diversity. The five PCs together explained 70% of the variance within the set of properties. The PC scores calculated based on all open-class words were highly correlated with the original PC scores calculated based on nouns (*rs* > 0.54).

**Table S4. AIC for the mixed effects models with different order effects of the gradient bins on psycholinguistic properties activation during speech.**

|  | Comprehension task | | | |  | Production task | | | |
| --- | --- | --- | --- | --- | --- | --- | --- | --- | --- |
| Orders | 0th | 1st | 2nd | 3rd |  | 0th | 1st | 2nd | 3rd |
| Action | -585.27 | **-585.62** | -583.78 | -582.31 |  | -893.78 | -895.69 | **-897.66** | -896.14 |
| AoA | -979.45 | **-981.76** | -981.10 | -981.50 |  | -1233.09 | -1235.42 | -1233.45 | **-1235.87** |
| Arousal | -604.97 | -610.67 | -613.40 | **-614.97** |  | -768.06 | **-771.82** | -769.83 | -767.99 |
| Closed-class words | 261.66 | 255.75 | 257.75 | **252.04** |  | 96.32 | 77.86 | **67.17** | 68.93 |
| Concreteness | -554.85 | -583.71 | **-586.69** | -584.69 |  | -818.49 | -834.26 | -854.50 | **-863.46** |
| Frequency | -559.98 | -558.57 | **-572.88** | -571.15 |  | -714.20 | -713.95 | **-727.43** | -725.44 |
| Global coherence | 83.34 | 84.99 | 76.53 | **75.81** |  | -71.15 | -84.91 | **-99.05** | -98.18 |
| Local coherence | 124.08 | 125.11 | 108.85 | **105.23** |  | **-66.28** | -64.92 | -64.69 | -63.53 |
| Number of phonemes | -996.55 | -996.48 | -1005.04 | **-1012.32** |  | -1179.77 | **-1191.35** | -1189.57 | -1187.60 |
| Number of words | -1448.41 | -1457.13 | -1463.76 | **-1496.69** |  | -1515.74 | -1516.68 | -1515.69 | **-1542.54** |
| Perception | -675.32 | **-691.64** | -690.12 | -689.92 |  | -831.07 | -829.72 | -837.85 | **-840.02** |
| SemD | -39.41 | -61.91 | -59.93 | **-63.30** |  | -251.94 | -249.97 | **-260.28** | -258.34 |
| Valence | -546.58 | -545.26 | -545.67 | **-547.91** |  | -756.72 | **-767.41** | -767.33 | -765.55 |

Note: AIC that suggested the best model fit for each property has been highlighted in bold.
